# Single-Cell Virtual Cytometer allows user-friendly and versatile analysis and visualization of multimodal single cell RNAseq datasets

**DOI:** 10.1101/843946

**Authors:** Frédéric Pont, Marie Tosolini, Qing Gao, Marion Perrier, Miguel Madrid-Mencía, Tse Shun Huang, Pierre Neuvial, Maha Ayyoub, Kristopher Nazor, Jean Jacques Fournié

**Author notes:** these authors contributed equally to this work (Frédéric Pont, Marie Tosolini).

## Abstract

The development of single cell transcriptomic technologies yields large datasets comprising multimodal informations such as transcriptomes and immunophenotypes. Currently however, there is no software to easily and simultaneously analyze both types of data. Here, we introduce Single-Cell Virtual Cytometer, an open-source software for flow cytometry-like visualization and exploration of multi-omics single cell datasets. Using an original CITE-seq dataset of PBMC from an healthy donor, we illustrate its use for the integrated analysis of transcriptomes and phenotypes of functional maturation in peripheral T lymphocytes from healthy donors. So this free and open-source algorithm constitutes a unique resource for biologists seeking for a user-friendly analytic tool for multimodal single cell datasets.

## INTRODUCTION

The recent development of techniques for single cell RNA sequencing (scRNAseq) has resulted in an accrual of scRNAseq datasets comprising thousands of cells from many lineages, tissues, physiological conditions and species. The classical representation of such datasets is based on their dimensionality reduction e.g. by t-stochastic neighborhood embedding (t-SNE) or uniform manifold approximation and projection (UMAP). In such steps, all cells are plotted according to their transcriptomic similarity with immediate neighbours and the overall community of cells, forming separate groups or clusters. The lineage, status and hallmarks of such cells and clusters are then identified by their expression levels of single genes, chosen for their hallmark expression patterns, *e.g.* expression of the *CD14* gene for monocytes, or of the *MKI67* gene for proliferating cells. Nevertheless, both technical noise from the data acquisition process, and massive gene dropouts impair detection of many genes in scRNAseq datasets. Consequently, mapping the expression level of a single gene in a t-SNE map is generally less informative than mapping the enrichment of a corresponding multi-gene signature.

We recently developed Single-Cell Signature Explorer, a tool which scores gene signatures by their UMI to total cell UMI ratio in each single cell from large datasets (1). This tool further overlays UMAP or t-SNE maps of the dataset with heatmap-encoded single cell scores of the signature. By allowing the user to see how these scores vary across all cells, it provides a visualization of any transcriptomic hallmark in the dataset. For example, this tool returns scores interpreted by user for identifying B versus non-B cells among peripheral blood mononuclear cells (PBMC). It may likewise help to infer single cell lineages, cell hallmarks, or any metabolic and proliferative status from collective gene expression levels (1).

However, formal identification of most cell lineages relies upon cell surface expression of canonical protein markers rather than on transcriptome-based inference. This issue was addressed by incepting the Cellular Indexing of Transcriptomes and Epitopes by Sequencing (CITEseq) (2), which allows simultaneous detection of single cell transcriptomes and antibody-labelled surface markers, yielding both gene expression levels and immunophenotypes from the same experiment. Several algorithms allow the analysis of either multidimensional immunophenotypings, *e.g.* Cytobank (3) or of single cell transcriptomes such as Seurat (4), iCellR, or Loupe Cell Browser (10×Genomics). However so far, there are no tools for the simultaneous and integrated visualization and analysis of both of these different data.

Here we introduce Single-Cell Virtual Cytometer to visualize both cell transcriptomes and phenotypes in t-SNE or UMAP from multimodal single cell datasets, as well as for flow-cytometry-like gating, selection and exploration of subsets of cells. We examplify its use to characterize the gene signatures for stages of functional maturation of peripheral T lymphocytes using CITEseq datasets of PBMC from healthy individuals. Our tool, implemented as freely available open-source software, represents a unvaluable resource to fully exploit the expanding universe of multi-omics datasets necessary to cancer research and care.

## MATERIALS AND METHODS

### Single-Cell Virtual Cytometer

Single-Cell Virtual Cytometer is a new tool, part of Single-Cell Signature Explorer software package (1) dedicated to high throughput signature exploration in single-cell analysis. It brings the flow cytometry software capabilities to single cell analysis. It is able to define and gate cell populations based on the 2D plot of, for example two antibodies or genes, and to display simultaneously the selected cells on a UMAP or t-SNE map. There is no limit to the number of antibodies, genes or other criteria possibly used to define the plot, including combinations of transcriptomic, proteomic and signature scores (1). Single-Cell Virtual Cytometer takes a data table as input, using tab-separated text file format, with the cells tag in rows and genes expression, antibodies detection levels, signature scores in columns and at least two columns with x,y coordinates for a map such as t-SNE or UMAP. Once the data table loaded, the user can then select two criteria, such as two antibodies. With such criteria, a flow cytometry-like contour plot of the entire dataset is drawn. Using a lasso or a box selection tool, the user can select some cells and immediately see these on a t-SNE/UMAP map. Single-Cell Virtual Cytometer supports an unlimited level of successive gatings. Selected cells can be exported for further use. Quadrant gates display the number and % of cells in each quadrant, and trigger their location on the t-SNE/UMAP.

### CITE-seq counter

We developed CITE-seq-counter software to count the UMI of antibodies tags in raw sequencing reads. CITE-seq-counter has been developed for Single Cell CITE-seq samples processed with 10XGenomics technologies. It takes as input fastq files R1 and R2 from the sequencer, the antibodies barcodes, and a white list of cells obtained from Seurat (4). Cell and antibody barcode positions are adjustable as well as UMI positions. Since sequencers can produce errors, one mismatch is allowed in the barcode and in the UMI. PCR duplicates (same UMI + barcode) are excluded from the counts. The software is written in Go, it is fast and the memory usage is as low as possible. Only the result table and two sequences R1 R2 are stored in RAM at the same time.

### Code availability

Single-Cell Virtual Cytometer was developed in pure javascript using the graphical libraries plotly.js (5) and Bulma. It only needs a web browser with javascript enabled to be executed, with tab-separated text files as input. Files can be accessed on the GitHub Single-Cell Virtual Cytometer web page.

CITE-seq-counter was developed in Go and pre-compiled static binaries are available for Linux and Windows.

### Generation and Preprocessing of PBMC CITE-seq data

Procedures for cell isolation, labeling, CITE-seq experiment, sequencing, and preprocessing of the resulting dataset are described in Supplementary Information section.

## RESULT

### Single-Cell Virtual Cytometer for analysis of CITE-seq datasets

Currently, there is a growing demand for analytic tools for exploration of CITE-seq datasets based on flow cytometry-like visualization of the cells. Thus we designed Single-Cell Virtual Cytometer, a software to explore scRNAseq datasets with user-friendly and flow cytometry-like tools. It can be run immediately in a web browser without requiring any complex installation. The user can immediately explore his data without mastering command line instructions. The data must consist in csv tables with cells in row and columns for genes, antibodies, signature scores, or any other quantitative single cell readout. Importantly, this table must have for each cell at least one set of map (x,y) coordinates from any dimensionality reduction method. Single-Cell Virtual Cytometer typically displays both a flow cytometry-like density plot of cell phenotypes (left panel) and the corresponding dimensionally-reduced map of cells based on their transcriptomes (right panel). Based on any user-defined criteria, the cells selected on the phenotype density plot by gates or quadrants are interactively displayed on the corresponding t-SNE /UMAP (Supplementary data demo video). Setting quadrants in the phenotype panel automatically triggers display of both percentages and counts of cells from each quadrant. Hence from a CITE-seq dataset analyzed with Single-Cell Virtual Cytometer, it is possible to select two antibodies to get their density plot across the cell population. Using gates or quadrants from this plot, the user can select further a subset of cells to visualize on the transcriptomic map. The cell tags of any selected subset of cells can be exported as a txt file. Downstream sub-gating and analysis with other antibodies of selected cells can be reiterated without limits. Delineating quadrants on the plot returns both the % and absolute count of cells from each quadrant, as well as their respective localization on the corresponding right side map. The plots and maps from Single-Cell Virtual Cytometer can be exported in low and high resolution. Importantly, the Single-Cell Virtual Cytometer is very versatile. Its two-panels displays are interactive and based on any (x,y) parameters selected by the user from the drop-down list. Hence this enables users to select not only any mAb or cell hashtag (Biolegend), but also any other parameter such as cluster number, sample annotation index, or dimensionality reduction axis. Hence instead of the above-depicted phenotype left and transcriptome right panels, the selection of (tSNE-1, tSNE-2) as left panel parameters allows user to gate, quadrant, and select cells from a transcriptomic cluster standpoint to further visualize their respective phenotypes on the right panel.

### Comparison with existing scRNAseq and flow cytometry visualization tools

#### Seurat 3.0

(4) is an R package designed for QC preprocessing, analysis, and exploration of single cell RNA-seq data. Seurat enables users to identify and interpret sources of heterogeneity from single-cell transcriptomic measurements, and to integrate diverse types of single-cell data, performing the so-called multimodal integration. Seurat is a command line tool able to generate dimensionality reduction maps from t-SNE or UMAP, allowing users to select clusters and subsets of cells using thresholds using command lines. Hence the use of Seurat requires bioinformatics skills that are however not necessary for using Single-Cell Virtual Cytometer, which was rather designed for end users more accustomed to flow cytometry.

#### Loupe Cell browser

3.1.1 is a dedicated visualization and analysis tool for scRNAseq developed for analysing scRNAseq datasets mostly produced by 10xGenomic platforms. It allows importing datasets and visualizing custom projections of either gene expression or antibody-only datasets,across t-SNE or UMAP computed by the Cell Ranger 3.1 pipeline. Despite its ease of use however, this tool only displays a single dimensionality reduced map featuring the dataset clusters and heatmaps of the graph-based differentially expressed genes or mAbs. Although this tool may export images and selection of cells, it lacks the dual displays of phenotypes and transcriptomes to perform any simultaneous exploration of CITEseq data.

### iCellR

is a R package for scRNAseq analysis able to produce interactive graphs for either of transcriptome or immunophenotype data, but not both simultaneously. To our knowledge iCellR does not reproduce a flow cytometry interface and, similarly to Seurat, is accessible for users skilled in R.

### CytoBank

(3) and some other flow cytometry softwares can import single cell data files after adequate file conversions, and can be used for visualizing single cell phenotype data. CytoBank is also able to produce t-SNE but its limited capacity to process a maximum of 818 parameters is not compatible with current scRNAseq transcriptomic datasets. Furthermore, Cytobank does not display interactively the density plot subpopulations in the t-SNE, nor does it allow users to export the cell tags for further use.

So currently, there are no other software than Single-Cell Virtual Cytometer to analyze CITE-seq or related multimodal single cell datasets as easily as with flow cytometry softwares. In addition, the computing time to display the user-selected plots and maps is extremely short. For example the time to plot a density plot or to display a mAb-labelled subpopulation on a t-SNE map is < 1sec with 10k cells when running Single-Cell Virtual Cytometer with an Intel(R) Xeon(R) CPU E5-2630 v4 @ 2.20GHz.

### Simultaneous visualisation of single cell phenotypes and transcriptomes from an healthy donor’s PBMC CITEseq dataset

Single-Cell Virtual Cytometer was primarily applied to analyse an original CITE-seq dataset of human PBMC. So the PBMC from a healthy individual were primarily labelled with a mix of 12 TotalSeqTM-B ADT (**Supplementary Table 1**) at 5 concentrations respectively labelled by 5 HTO (**Supplementary Table 2**). The stained PBMC were analyzed for CITE-seq using a 10XGenomics 3’ chemistry V3 platform, sequenced, pre-processed, and dimensionality reduction of the transcriptome datasets was performed with UMAP. As QC of the transcriptomic part of the CITE-seq dataset, the single cells with outlier counts of total UMI or number of genes were discarded. For the QC of the phenotypic part of the CITE-seq dataset, cells labelled with over-diluted ADT (**Supplementary Figure 1**) and cells displaying mutually exclusive phenotypes (e.g. CD3^+^CD19^+^CD14^+^) were discarded. This finally yielded a CITE-seq dataset encompassing the phenotype and transcriptome of n= 5,559 PBMC. Importantly, analysing this dataset by either the flow cytometry tool Cytobank or by Single-Cell Virtual Cytometer yielded the same plots and rates of CD3CD4 T and CD3CD8 T cells from the PBMC (**Figure 1**).

**Figure 1.**
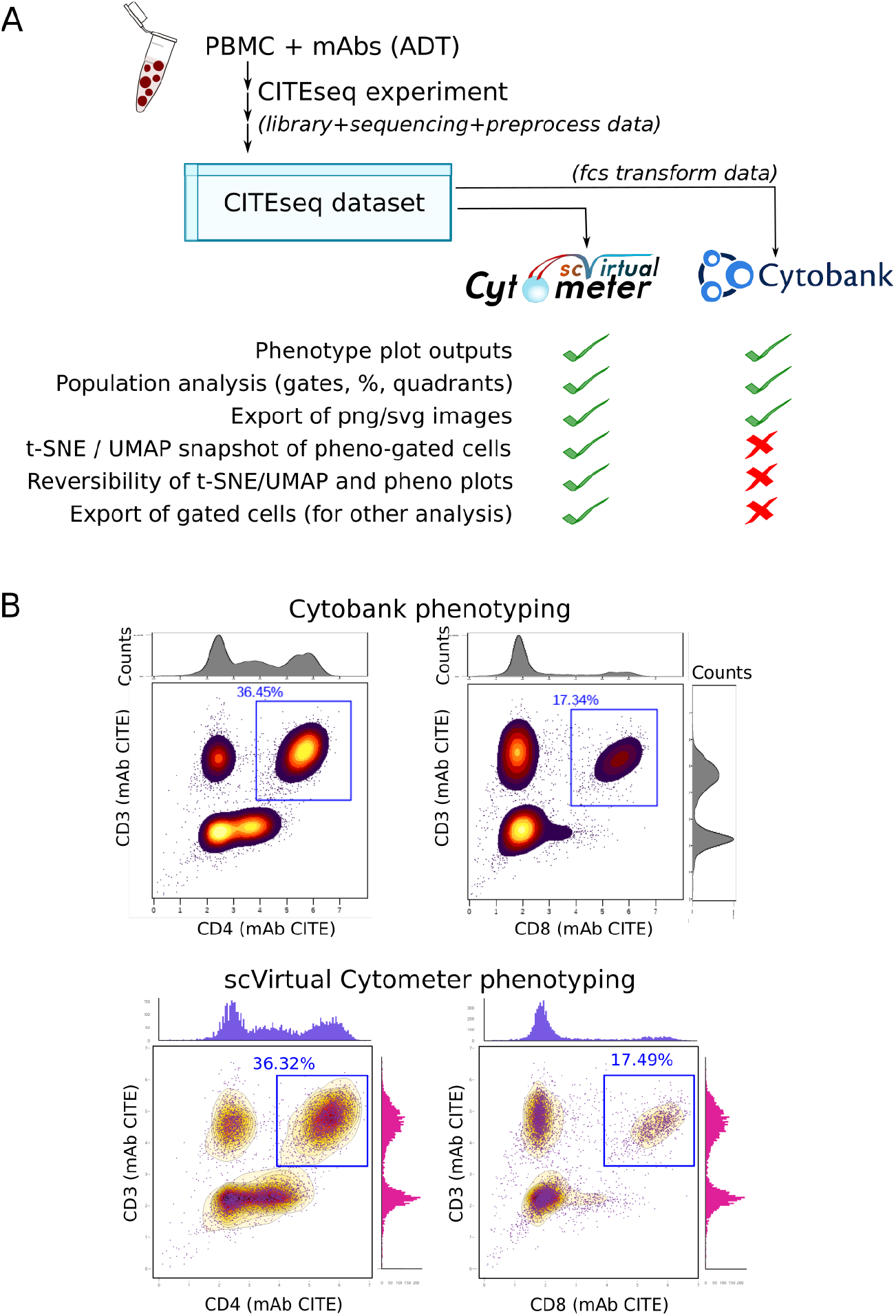
A: The PBMC from a healthy individual were labeled with ADT and studied by CITE-seq prior to analysis of the cell phenotype results by either the flow cytometry software Cytobank or by Single-Cell Virtual Cytometer. B: Examples of phenotype analysis by either software, illustrating the consistence of their displays and results.

We reasoned that the above-depicted possibility to explore at the same time phenotype data and single gene expressions opens the possibility to explore likewise phenotype data and multi-gene signatures. Any multi-gene signature can be robustly scored for each single cell across the scRNAseq dataset by computing its summed expression ratio to that cell’s transcriptome (1). So the above CITEseq PBMC dataset from one donor was then analysed in more details through both gene signatures and cell surface markers, by computing (Methods) and plotting the single cell scores of a myeloid cell-specific gene signature (1) *versus* ADT staining for the T cell surface marker CD3. The T lymphocytes were then gated and visualized across the UMAP of the entire dataset (**Figure 2A**). These gated T cells were further analyzed likewise for cell surface CD4 *versus* CD8 markers, which yielded the subsets of CD4 T (*n*=1796), CD8 T (*n*=885), CD4CD8 double positive T cells (*n*=52) and the CD4CD8 double negative T cells (*n*=99) including *γδ* T lymphocytes (6). These four subsets were readily delineated in the dimension-reduced UMAP of the PBMC dataset (**Figure 2B**). Parallel analyses of CD19 and CD16 phenotypes in the non-T cell subsets of PBMC identified the B lymphocytes (CD19^+^CD16^−^) (*n*=157), the NK cells (CD19^−^CD16^+^) (*n*=902) and monocytes (CD19−CD16−) (*n*^=^1341), including their classic CD14^+^CD16^−^ and non-classic CD14^*int*^CD16^*int/*+^ subsets (**Figure 2C**).

**Figure 2.**
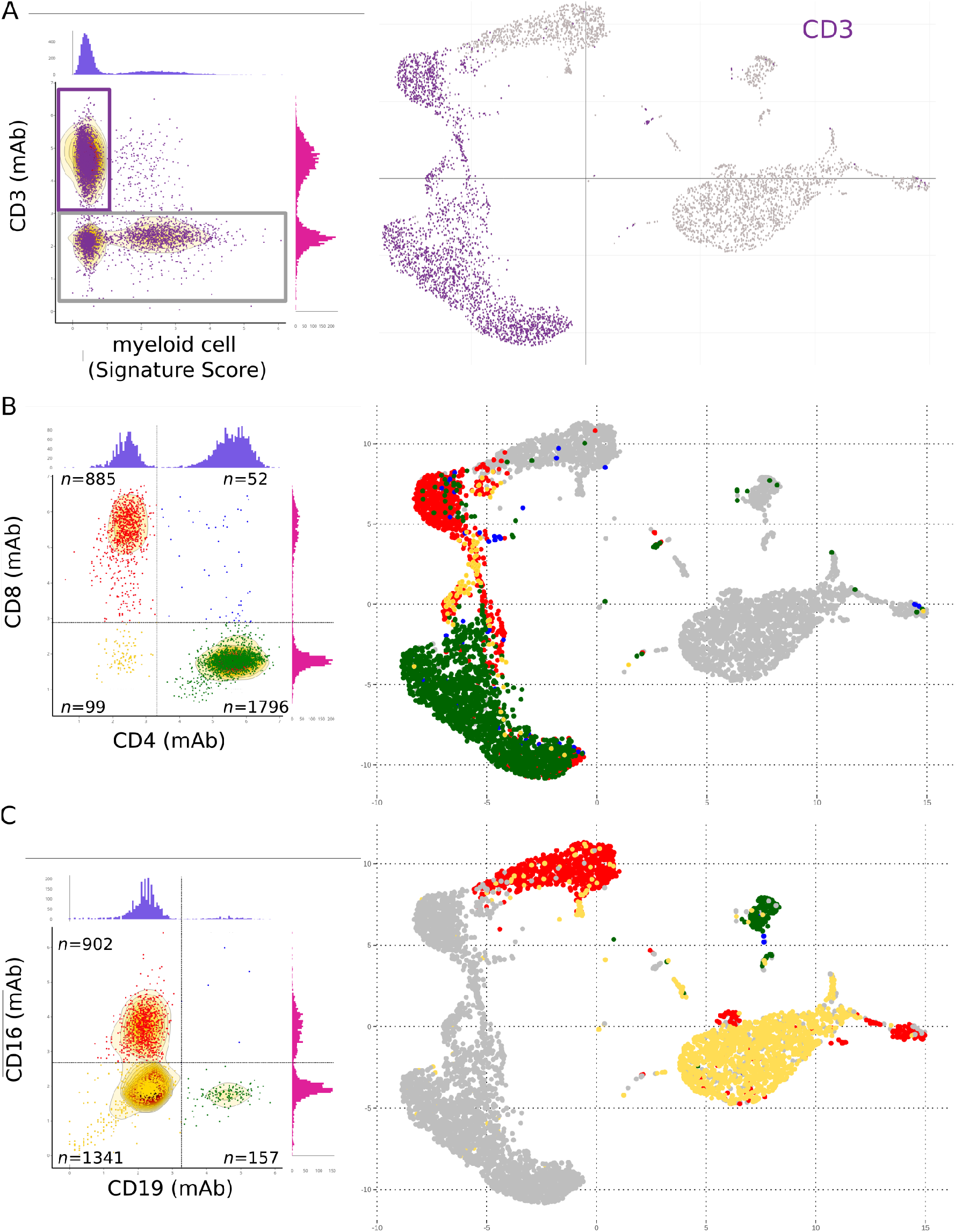
Simultaneous visualization by Single-Cell Virtual Cytometer of cell surface phenotype, gene signatures and cell subsets in the UMAP of 6k PBMC isolated from an healthy individual and stained with TotalSeq-A^TM^ADT. A: *Left panel:* The scores for a myeloid gene signature (*CD14, LYZ, ANPEP, FUT4, S100A2,S100A4-S100A6, S100A8-S100A13, S100B* genes) *versus* CD3 staining levels of 6k PBMC define the T lymphocytes (purple gate) further shown in the transcriptome UMAP *(right panel)*. B: The CD4 and CD8 phenotype of the above-gated T lymphocytes *(left panel)* defines four T cell subsets shown in the corresponding transcriptome UMAP *(right panel)*. C: CD19 and CD16 phenotype of non-T cells from PBMC *(left panel)* defines the B, NK, and myeloid cell subsets respectively shown in the corresponding transcriptome UMAP *(right panel)*.

So, Single-Cell Virtual Cytometer allows the simultaneous analysis and visualization of cell surface transcriptomes and phenotypes at the single cell level.

### Consistence of gene signatures and cell surface phenotype of peripheral T cell differentiation at the single cell level

Peripheral T lymphocytes evolve through successive differentiation stages comprising naive (Tn), central memory (Tcm), effector memory (Tem) and terminally differentiated (Temra) lymphocytes (7). These respective stages are classically defined through cell surface expression of markers such as CD45RA and CD62L, IL7Ra, and CCR7 (Tn), CD45RO, CD62L, IL7Ra and CCR7 (Tcm), CD45RO (Tem), and CD45RA (Temra) proteins, but their respective transcriptome signatures have remained unclear as cell surface markers and transcriptomes have never been studied on the same cells so far.

In a test experiment with the above CITE-seq dataset and Single-Cell Virtual Cytometer, we now aimed at defining these differentiation signatures. The cell surface expression of CD4 and CD8 markers indicated that T lymphocytes comprised *n*= 1796 CD4 T cells, *n*= 885 CD8 T cells, *n*= 52 CD4CD8 double positive T lymphocytes, and *n*= 99 double negative T lymphocytes, embedded in distinct areas of the PBMC dataset UMAP (**Figure 2B**). These four subsets of T lymphocytes were then subdivided within Tn, Tcm, Tem, and Temra based on their cell surface CD45RA and CD62L phenotypes **(Figure 3)**. In CD4 T cells, this identified CD45RA and CD62L-double positive cells corresponding to CD4 Tn lymphocytes, CD45RA-negative CD62L-positive cells corresponding to CD4 Tcm lymphocytes, *n*= 438 CD45RA and CD62L-double negative cells corresponding to CD4 Tem lymphocytes, and *n*= 10 cells that were both CD45RA-positive and CD62L-negative, corresponding to the CD4 Temra lymphocytes. The genes selectively and differentially up-regulated by Tcm *versus* Tn cells, by Tem *versus* Tcm cells, and by Temra *versus* Tem cells, and by Tn *versus* all other cells were selected (BH-corrected Wilcoxon *P* < 0.001). This defined four differentiation signatures which were refined by discarding genes with intra-group mean <0.1 and relative variance >1. These differentiation signatures were then scored across each single cell of CD4 T lymphocyte, and the same procedure was applied separately for the differentiation signatures of CD8 T lymphocytes, double positive T lymphocytes, and double negative T lymphocytes **(Supplementary Tables 3-6)**. For each of these gated T cell subsets, these differentiation signatures were consistent with the corresponding cell surface CD45RA and CD62L phenotype **(Figure 3)**.

**Figure 3.**
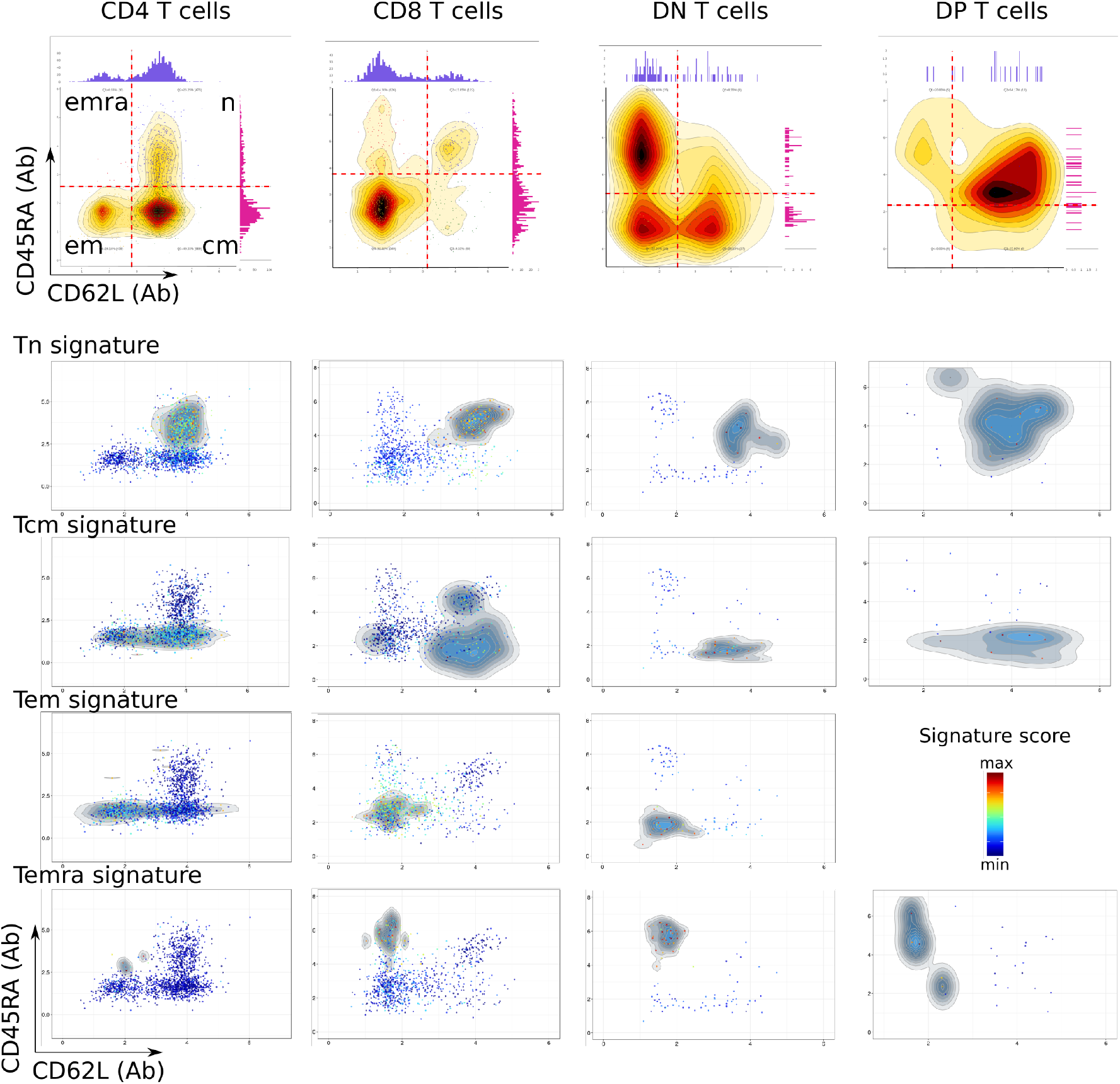
Cell surface phenotype (top) and gene signatures (bottom) of differentiation stages in T lymphocytes among 6k PBMC from an healthy individual. The gated CD3-positive cells were subdivided according to cell surface markers as CD4 T, CD8 T, double negative T, and double positive T lymphocytes. Each of these subset was then gated and analyzed for expression of the cell surface CD62L and CD45RA markers. This dataset did not encompass Tem cells among the DP T lymphocytes.

In a further validation experiment, the T cells from another PBMC CITE-seq dataset were analyzed with Single-Cell Virtual Cytometer as above. The 10k PBMC CITE-seq (3’ chemistry V3) dataset was downloaded from the 10XGenomics website, pre-processed with Seurat as above. The CD4, CD8, DN, and DP T lymphocytes were identified by their CD3, CD4, and CD8 phenotype, and scored for the above differentiation signatures. Within this second PBMC dataset, all the transcriptome signatures of T cell differentiation were also consistent with the differentiation phenotypes, here defined by the cell surface expression of IL7R (CD127) and CD45RA **(Supplementary Figure 2)**.

Together, these results validated the simultaneous analysis and visualization of gene expression and cell surface phenotype from CITE-seq datasets with Single-Cell Virtual Cytometer.

## DISCUSSION

The advent of Cellular Indexing of Transcriptome and Epitopes by sequencing (CITE-seq) has brought the possibility to jointly obtain both gene expression and immunophenotypes at the single cell level (2). However, this development does not provide a user-friendly interface allowing biologists to readily visualize any combination of gene, immunophenotype marker and combination signature from the datasets.

Our report illustrates the potential of Single-Cell Virtual Cytometer for the exploration of both transcriptomic and phenotype from CITE-seq datasets. This simple and rapid software does not intervene on the pre-processing integration, clusterization and QC of data however, in contrast to pre-existing algorithms and platforms that analyze either of phenotypes or transcriptomes separately. This unique tool is particularly relevant for fully exploiting approaches that measure distinct modalities within single cells, since each readout such as gene, protein or signature score is directly plotted across the dataset plot. By pinpointing here the gene signatures of functional differentiation stages in peripheral T lymphocytes from healthy individuals, we showed that it permits straightforward analyses of bimodal data such as mRNA and cell surface proteins.

Likewise, Single-Cell Virtual Cytometer is broadly applicable to the visualization of any kind of readout from any multimodal single cell technology, after adequate integration of the data sets (8). Its versatility enables users to analyze likewise any kind of single cell data about chromatin accessibility (9, 10), epigenomics (11), mutations (12), chromosome conformation (13), RNA modification (14), spatial transcriptomics (15, 16) and spatial proteomics (17, 18). Hence, Single-Cell Virtual Cytometer represents a unvaluable resource for integrated analyses of multimodal datasets at the single cell level.

## ACKNOWLEDGEMENTS

This work was supported by institutional grants from CNRS (SCALES project from Mission “Osezx l’interdisciplinarité”) This work was granted access to the HPC resources of CALMIP supercomputing center under the allocation 2019-T19001. We are also grateful to the Génotoul bioinformatics platform (Bioinfo Genotoul, Toulouse Midi-Pyreneés) for providing computing resources.

## Conflict of interest statement

Qing Gao,Tse Shun Huang and Kristopher Nazor are employees of Biolegend.

