## Supplementary Information for "Single-Cell Virtual Cytometer allows user-friendly and versatile analysis and visualization of multimodal single cell RNAseq datasets"

### Supplementary figures

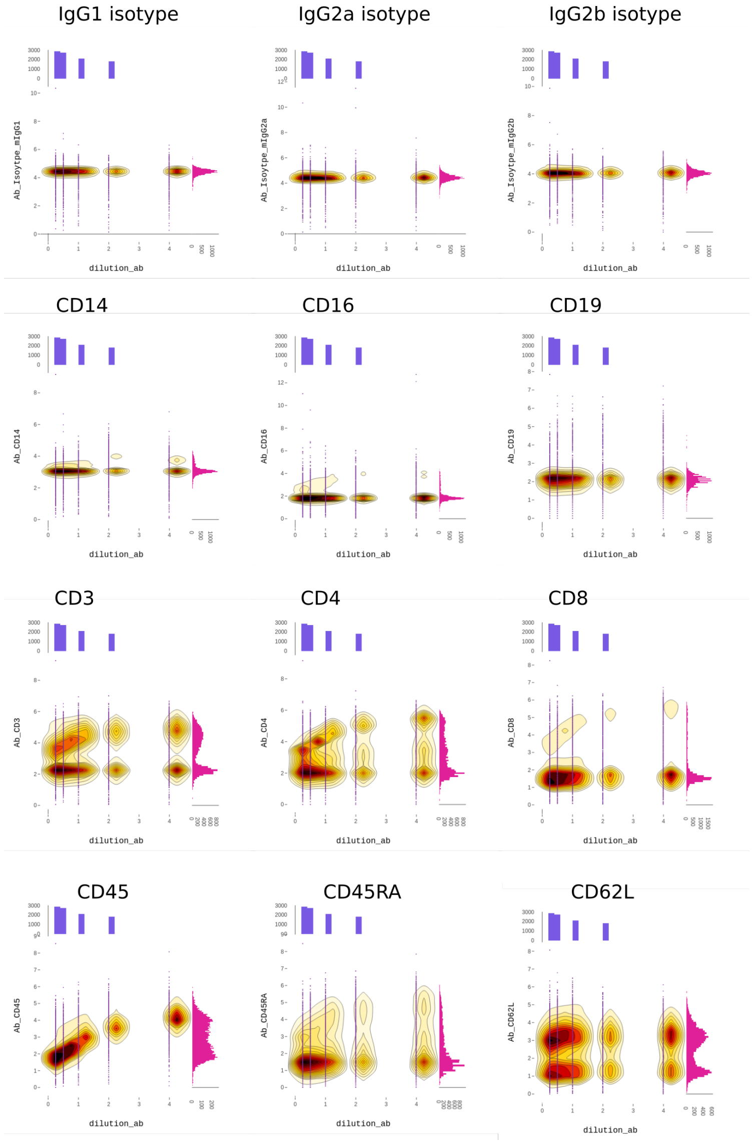


**Supplementary Figure 1.** Visualization by Single-Cell Virtual Cytometer of each ADT titrated by HTO in the CITE-seq data set of 8k PBMC from an healthy individual. For each specified antibody, all dataset cells are plotted for ADT dilution (x axis) *versus* intensity of antibody staining (y axis). The associated purple and pink histograms show the corresponding density distributions of x and y parameters, respectively.

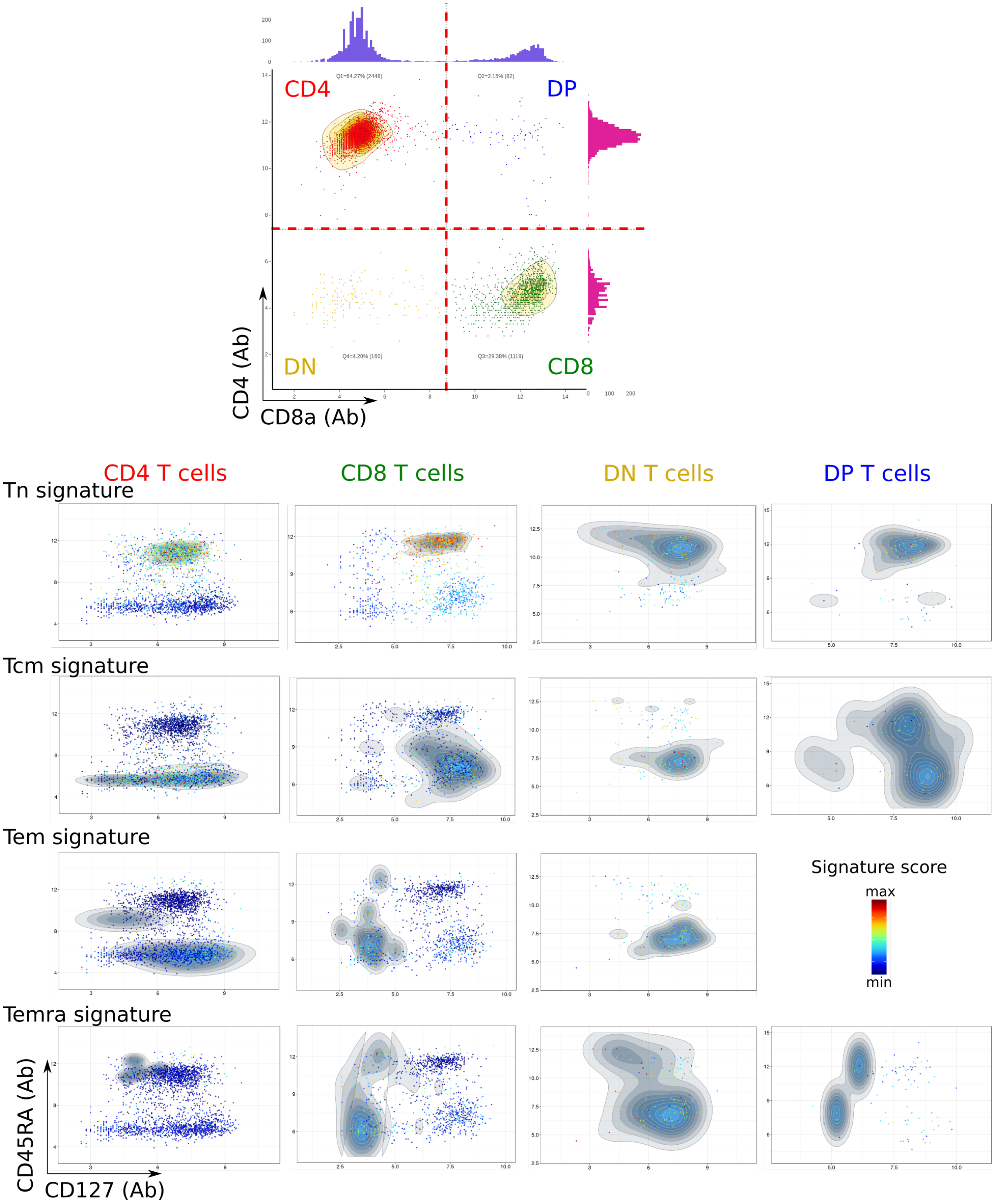


**Supplementary Figure 2.** The CD4, CD8, DN, and DP T cells from a validation CITE-seq 10XGenomics dataset of 10k PBMC stained with TotalSeq-B$^{TM}$ADT were defined and gated using the CD3, CD4, and CD8a antibodies (top), and then analyzed in parallel for their respective pattern of expression of the cell surface differentiation markers IL7R (CD127) and CD45RA (bottom). Heatmap shows the scores for the same differentiation signatures as in Figure 3. The signatures are listed in Supplementary Tables 3-6.

### Supplementary Material and methods

#### Single-Cell Signature Scorer

Single-Cell Signature Scorer[1] was used to calculate geneset enrichment scores for single cell transcriptomes. Briefly, scRNAseq data were processed using Seurat 3.0 toolkit package[2] involving the normalization and variance stabilization package sctransform[3]. Then, scores were computed for each single cell as described in[1] using genesets from MSigDB[4,5] as well as additional user-defined genesets (list of HUGO gene symbols in a text file format, Supplementary Tables 3-6).

#### Isolation of PBMC

Human blood from healthy adults were collected with informed consent of the donor in sodium heparin tubes (Becton Dickinson), and the peripheral blood mononuclear cells (PBMCs) were isolated by sedimentation on Lymphopure™(BioLegend).

#### Cell surface labelling of PBMC

Five TotalSeq™-A antibody mixes (Supplementary Table 1) were used for labeling of PBMCs with a five-point titration curve (4X, 2X, 1X, 0.5X, 0.25X, Supplementary Table 2). For each antibody mix, a different TotalSeq™-A hashtag (0.5$\mu$g/test) was used to allow for sample multiplexing (Supplementary Table 2). For each titration sample, 1 million PBMC were blocked with Human TruStain FcX™(BioLegend) using 5 $\mu$L brought to a final volume of 50 $\mu$L with PBS at 4°C for 10 min in 5 mL polypropylene round-bottom tubes (Falcon). Cells were then incubated with the antibody mixtures at 4°C for 30 min, followed by three washes with 3.5 mL of Cell Staining Buffer (BioLegend). After the last wash, samples were resuspended in 500$\mu$L Cell Staining Buffer (BioLegend) to get an approximate concentration of 2x$10^6$ cells/mL, and 40 $\mu$m Flowmi™Cell Strainer (Bel-Art, H-B Instruments) were used to remove cell clumps. To ensure that each sample was appropriately mixed into the same 10X Chromium lane, its cell concentration and viability were individually adjusted to $10^6$ cells/mL using the Countess™II FL Automated Cell Counter, added in equal volume to the mixture, and the final concentration of the mixed hashed sample was verified by Cell Counter.

#### CITE-seq RNA and Antibody library generation

Libraries containing mRNAs, antibody derived tags (ADTs) and hashtag oligonucleotides (HTOs) were generated using Single-Cell 3′ chemistry V3 kit (10X Genomics) according to the manufacturer's instructions (Document CG00018, RevB). A total of approximately 40,000 cells from the final mixture was loaded in one 10X lane, for gel bead-in-emulsion (GEM) generation and barcoding, with an expected cell recovery of 25,000 cells corresponding to 5000 cells per titration sample. To accommodate the generation of TotalSeq™-A ADT/HTO libraries, the following modifications were made. In step 2.2, the 35 $\mu$L mixture containing cDNA, ADTs and HTOs were mixed with 50 $\mu$L Amp Mix (Chromium Single Cell 3′ GEM Module), 15 $\mu$L Feature cDNA primers (Chromium Single Cell 3′ GEM Module), 1 $\mu$L ADT additive primer (0.2 $\mu$M stock, 5′CCTTGGCACCCGAGAATT*C*C), 1 $\mu$L HTO additive primer (0.2 $\mu$M stock, 5′GTGACTGGAGTTCAGACGTGTGC*T*C), followed by PCR amplification as described in the protocol. The PCR conditions were: an initial denaturation step at 98°C for 3 min, followed by 12 cycles of denaturation at 98°C for 15 sec, annealing at 63°C for 20 sec, and extension at 72°C for 1 min, followed by final extension at 72°C for 1 min. cDNA, ADTs and HTOs were separated and purified after amplification by following Steps 2.3A and 2.3B from manufacturer's Document CG00018, RevB. After Step 2.3B from CG00018, RevB, the ADT libraries were PCR-amplified using 50 $\mu$L Quantabio sparQ HiFi PCR Master Mix (Quantabio), 40 $\mu$L Nuclease-free Water (Thermo Fisher), 2.5 $\mu$L SI-PCR primer (5′AATGATACGGCGACCACCGAGATCTACACTCTTT-CCCTACACGACGC*T*C) at 10$\mu$M, 2.5 $\mu$L RPI3 (10 $\mu$M stock, TruSeq Small RNA RPIx, Illumina) and 5 $\mu$L end product from Step 2.3B. PCR conditions were: 1 cycle at 98°C for 2 min, followed by 13 cycles at 98°C for 20 sec, 60°C for 30 sec, and 72°C for 20 sec, followed by 1 cycle at 72°C for 5 min. HTO libraries were generated by PCR using 50 $\mu$L Quantabio sparQ HiFi PCR Master Mix, 40 $\mu$L Nuclease-free Water, 2.5 $\mu$L SI-PCR primer (10 $\mu$M stock), 2.5 $\mu$L D703-s (10 $\mu$M stock, TruSeq D70x-s, Illumina) and 5 $\mu$L end product from Step 2.3B (Manufacturer's document CG00018, RevB). PCR conditions were: 1 cycle at 98°C for 2 min, followed by 11 cycles at 98°C for 20 sec, 64°C for 30 sec, and 72°C for 20 sec, followed by 1 cycle at 72°C for 5 min. ADT/HTO libraries were purified and quantified following Step 4.2 to Step 4.3 of manufacturer's document CG00018, RevB.

#### Sequencing

ADT, HTO, and RNA libraries were sequenced on an Illumina NovaSeq6000 platform with a NovaSeq 6000 S1 Reagent Kit, 300 cycles (Illumina) to a read depth of 25,000 reads/cell for mRNA, 5000 reads/cell for ADTs, 500 reads/cell for HTOs. Sequencing parameters were set for Read1 (151 cycles), Index1 (8 cycles), and Read2 (151 cycles). When trimmed to 91 bp the read2 yielded cellranger counts almost similar and correlated (Median $r > 0.95$ (Spearman) and $>0.99$ (Pearson)) to that of 150bp read2, indicating that length of RNA read 2 did not affect the identification and expression profiles of single cells.

#### Preprocessing of CITE-seq data

The CITE-seq RNA reads were mapped to the human genome (GRCh38) and transcripts were quantified as indicated above. Antibody counts for CITE-seq[6] were counted using CITE-seq-counter (see above) for the 10XGenomics chemistry V2 datasets, and with Cell Ranger 3.0 for the 10XGenomics chemistry V3 datasets.The result is a table with cells in rows and antibodies in columns, which is further merged with the transcriptomic data and XY map coordinates using Single-Cell Signature Merger[1]. The t-SNE or UMAP plots were produced by Seurat[2] as follows: the raw data (fastq files) were computed with CellRanger 3.0 and then loaded in a R session with the Seurat 3.0 toolkit package involving the normalization and variance stabilization package sctransform[3]. Samples were individually filtered using UMI and percentage of mitochondrial genes criteria. Samples were then merged using batch correction to align datasets as described[2]. t-SNE or UMAP coordinates were then calculated using the 11 first PCA and exported in a table. The PBMC CITE-seq dataset generated in this study can be downloaded from (Gene Expression Omnibus, GSE number pending).

### Supplementary Tables

| Marker | Clone | microgram per test(1x) | Category | Barcode | Barcode sequence | Ref. | Vendor |
|---|---|---|---|---|---|---|---|
| Mouse IgG2b, kappa isotype Ctrl | MPC-11 | 1.00 | TotalSeq$^{TM}$-A | 0092 | ATATGTATCACGCGA | 400373 | BioLegend |
| Mouse IgG1, kappa isotype Ctrl | MOPC-21 | 1.00 | TotalSeq$^{TM}$-A | 0090 | GCCGGACGACATTAA | 400199 | BioLegend |
| Mouse IgG2a, kappa isotype Ctrl | MOPC-173 | 1.00 | TotalSeq$^{TM}$-A | 0091 | CTCCTACCTAAACTG | 400285 | BioLegend |
| CD19 | HIB19 | 0.125 | TotalSeq$^{TM}$-A | 0050 | CTGGGCAATTACTCG | 302259 | BioLegend |
| CD3 | UCHT1 | 0.25 | TotalSeq$^{TM}$-A | 0034 | CTCATTGTAACTCCT | 300475 | BioLegend |
| CD16 | 3G8 | 0.50 | TotalSeq$^{TM}$-A | 0083 | AAGTTCACTCTTTGC | 302061 | BioLegend |
| CD4 | RPA-T4 | 0.125 | TotalSeq$^{TM}$-A | 0072 | TGTTCCCGCTCAACT | 300563 | BioLegend |
| CD8 | SK1 | 0.25 | TotalSeq$^{TM}$-A | 0046 | GCGCAACTTGATGAT | 344751 | BioLegend |
| CD14 | 63D3 | 1.00 | TotalSeq$^{TM}$-A | 0051 | CAATCAGACCTATGA | 367131 | BioLegend |
| CD45 | 2D1 | 0.125 | TotalSeq$^{TM}$-A | 0048 | TCCCTTGCGATTTAC | 368543 | BioLegend |
| CD45RA | HI100 | 0.10 | TotalSeq$^{TM}$-A | 0063 | TCAATCCTTCCGCTT | 304157 | BioLegend |
| CD62L | DREG-56 | 0.13 | TotalSeq$^{TM}$-A | 0147 | GTCCCTGCAACTTGA | 304847 | BioLegend |

**Supplementary Table 1.** Titration of Antibodies used in Figures 1 and 2

| mAb titration | Hashtag | Clone | microgram per test | Category | Barcode | Barcode sequence | Ref | Vendor |
|---|---|---|---|---|---|---|---|---|
| 0.25x | Anti-human hashtag 1 | LNH-94; 2M2 | 0.5 | TotalSeq$^{TM}$-A | 0251 | GTCAACTCTTTAGCG | 394601 | BioLegend |
| 0.50x | Anti-human hashtag 2 | LNH-94; 2M2 | 0.5 | TotalSeq$^{TM}$-A | 0252 | TGATGGCCTATTGGG | 394603 | BioLegend |
| 1.00x | Anti-human hashtag 3 | LNH-94; 2M2 | 0.5 | TotalSeq$^{TM}$-A | 0253 | TTCCGCCTCTCTTTG | 394605 | BioLegend |
| 2.00x | Anti-human hashtag 4 | LNH-94; 2M2 | 0.5 | TotalSeq$^{TM}$-A | 0254 | AGTAAGTTCAGCGTA | 394607 | BioLegend |
| 4.00x | Anti-human hashtag 5 | LNH-94; 2M2 | 0.5 | TotalSeq$^{TM}$-A | 0255 | AAGTATCGTTTCGCA | 394609 | BioLegend |

**Supplementary Table 2.** List of human TotalSeq$^{TM}$-A hashtags used for CITE-Seq of PBMC

| naive CD4 | CM CD4 | EM CD4 | EMRA CD4 |
|---|---|---|---|
| CHI3L2 | CTSA | STOM | TCHP |
| STMN1 | NPDC1 | MATK | RNF5 |
| ADTRP | ITGB1 | CGAS | HSPA1A |
| TMIGD2 | | PLCB1 | C2CD5 |
| BACH2 | | BHLHE40 | UBE2D4 |
| EPHA1.AS1 | | RHOC | STX7 |
| TTN | | TNFRSF18 | LMNB2 |
| ACTN1 | | MYBL1 | PDZD4 |
| EPHX2 | | KLRG1 | PER2 |
| APBA2 | | CYTOR | GPD1L |
| ICA1L | | HOPX | MPP1 |
| AK5 | | LINC01871 | LAIR2 |
| PDK1 | | LGALS3 | DOCK5 |
| CHRM3.AS2 | | LYAR | TGFBR3 |
| FHIT | | CST7 | TCF7L2 |
| AIF1 | | LINC01934 | AL031848.2 |
| CCR7 | | | ZW10 |
| IL6ST | | | PTCH1 |
| | | | SECTM1 |
| | | | FAM219A |
| | | | S1PR5 |
| | | | ALG14 |
| | | | ARRDC4 |
| | | | NAP1L2 |
| | | | RGS9 |
| | | | C1orf21 |
| | | | FBXO30 |
| | | | GPR137B |
| | | | ADGRG1 |
| | | | PLGLB1 |
| | | | TTYH3 |
| | | | PRSS23 |
| | | | CES1 |
| | | | BACE1 |
| | | | CEBPA |
| | | | CD1D |
| | | | MCTP1 |
| | | | SLC25A43 |
| | | | SPATA21 |
| | | | MYOF |
| | | | AC243829.1 |
| | | | ZEB2 |
| | | | CX3CR1 |
| | | | CPM |
| | | | KYNU |
| | | | SKAP2 |

**Supplementary Table 3.** Genes used in identification of CD4 step maturation

| naive CD8 | CM CD8 | EM CD8 | EMRA CD8 |
|---|---|---|---|
| KCNQ1OT1 | GPR183 | SYTL2 | KLRF1 |
| LINC02446 | SOCS3 | ASCL2 | CMC1 |
| TMEM204 | MAP3K1 | PRDM1 | LAIR2 |
| RGCC | CD28 | MT1E | CD160 |
| TTN | CD82 | ZNF683 | PDLIM1 |
| MYC | BNIP3 | MIR4435.2HG | CCL4L2 |
| ITGA6 | NPDC1 | THEMIS2 | KIR3DL2 |
| CLEC11A | CASK | ADRB2 | IKZF2 |
| RALGPS2 | | RHBDF2 | TRGV9 |
| LRRN3 | | CD226 | SYNGR1 |
| LMO7 | | GTF3C1 | AC090152.1 |
| BIRC3 | | PLEKHG3 | PTGDR |
| AC138150.1 | | FGFBP2 | CTSZ |
| PITPNA.AS1 | | CYTOR | CCL3L1 |
| RUNX2 | | | TIGIT |
| AC025164.1 | | | ARHGAP18 |
| EPHA1.AS1 | | | AOAH |
| AK5 | | | CEP78 |
| FAM102A | | | |
| RAB3GAP1 | | | |
| PIM2 | | | |
| LINC00402 | | | |
| TMEM63A | | | |
| BEX3 | | | |
| HIVEP2 | | | |
| ZBTB10 | | | |
| CFAP20 | | | |
| SLC7A6 | | | |
| PASK | | | |
| HSPB1 | | | |
| ETFRF1 | | | |
| AC136475.3 | | | |
| SNHG19 | | | |
| AL445686.2 | | | |
| LYRM4 | | | |
| CITED4 | | | |
| RGL4 | | | |
| LINC00476 | | | |
| CHRM3.AS2 | | | |
| MAL | | | |
| BSDC1 | | | |
| SYPL1 | | | |
| EPHX2 | | | |
| TMEM106B | | | |
| ICA1L | | | |
| TMEM14C | | | |
| GPR155 | | | |
| PRKCA | | | |
| DNMT3A | | | |
| CHMP7 | | | |
| AC097376.2 | | | |
| COA1 | | | |
| THEM4 | | | |
| TBC1D4 | | | |
| ZBTB25 | | | |
| ACTN1 | | | |
| EBPL | | | |
| CD55 | | | |
| BEX4 | | | |
| LINC01550 | | | |
| NCF1 | | | |
| DNPH1 | | | |
| A1BG | | | |
| DGKA | | | |
| ZNF609 | | | |
| RNF157 | | | |
| ZNF274 | | | |
| AL662844.4 | | | |
| ZEB1 | | | |
| SPINT2 | | | |
| IPCEF1 | | | |

*Continued on next page*

Supplementary Table 4 – *Continued from previous page*

| naive CD8 | CM CD8 | EM CD8 | EMRA CD8 |
|---|---|---|---|
| APBA2 | | | |
| BICDL1 | | | |
| FCGRT | | | |
| MLXIP | | | |
| YBX3 | | | |
| RETREG1 | | | |
| SREBF1 | | | |
| SH3YL1 | | | |
| ZNF101 | | | |
| EEF1G | | | |
| PDK1 | | | |
| INPP4B | | | |
| SELENOM | | | |
| PCED1B | | | |
| CRLF3 | | | |
| RIC3 | | | |
| TXK | | | |
| ATF7IP2 | | | |
| PDE7A | | | |
| TMIGD2 | | | |
| SGTB | | | |
| BEX2 | | | |
| IL6ST | | | |
| TCP11L2 | | | |
| NFKB1 | | | |
| ALG13 | | | |
| AL136454.1 | | | |
| CAMK4 | | | |
| PDE3B | | | |
| ITPKB | | | |
| MAML2 | | | |
| CCR7 | | | |
| AIF1 | | | |
| LDLRAP1 | | | |
| TRABD2A | | | |
| STMN3 | | | |
| OXNAD1 | | | |
| GABPB1.AS1 | | | |
| LEF1 | | | |
| NFKBIZ | | | |
| GIMAP1 | | | |
| ALKBH7 | | | |
| ATP6V0E2 | | | |
| MCUB | | | |
| SATB1 | | | |
| ABLIM1 | | | |
| NBEAL1 | | | |
| NOSIP | | | |
| PRMT2 | | | |
| TCF7 | | | |
| LEPROTL1 | | | |
| NUCB2 | | | |
| C6orf48 | | | |
| PIK3IP1 | | | |
| FOXP1 | | | |
| RGS10 | | | |

**Supplementary Table 4.** Genes used in identification of CD8 step maturation

| naive DN | CM DN | EM DN | EMRA DN |
|---|---|---|---|
| RPS17 | EMC7 | IL7R | FYN |
| GIMAP7 | FNIP1 | GBP5 | TRAPPC1 |
| DDX18 | GSPT1 | EIF4A1 | ATP8A1 |
| LDHB | HNRNPUL1 | ALOX5AP | AP2S1 |
| NEMF | TRGC1 | SRM | TRBC1 |
| ARID1B | PSMD2 | PRMT1 | LSM10 |
| TMEM123 | HBP1 | PRKRA | PAFAH1B1 |
| GDI2 | RAB11A | CDC26 | SLTM |
| STRAP | BSDC1 | CAMK4 | RPA2 |
| ALKBH5 | CTNNBL1 | PMVK | NDUFB5 |
| NUB1 | RAB5C | DRG2 | TULP4 |

*Continued on next page*

Supplementary Table 5 – *Continued from previous page*

| naive DN | CM DN | EM DN | EMRA DN |
|---|---|---|---|
| ADAR | PRKDC | OGFR | THOC2 |
| C6orf48 | HNRNPL | TC2N | CCDC14 |
| TCF7 | REV1 | PPIL2 | SLC2A4RG |
| ZC3HAV1 | API5 | NUDT21 | SREK1IP1 |
| SSBP1 | ITGAE | FAM98B | PET117 |
| MECP2 | TXNRD1 | C15orf61 | NGDN |
| GNB1 | HMG20B | E2F4 | CMC1 |
| MRPS34 | NFKB1 | SND1 | DOCK11 |
| DYRK2 | TSNAX | PMPCA | TFDP2 |
| RIOK3 | ELOA | CDC42EP3 | HLA.DPB1 |
| RBBP6 | SURF1 | NR1D2 | TRGC2 |
| LSM5 | PRDX2 | PTBP3 | NAE1 |
| SYS1 | PFDN4 | C19orf24 | CD320 |
| PARVG | MAP2K1 | TRADD | PITPNM1 |
| TRAPPC6A | QTRT1 | LSM6 | PPP1R18 |
| NGLY1 | COQ4 | GYG1 | ZMYM2 |
| RECQL | TTC5 | GOLGA3 | RAB6A |
| PER1 | MRPL33 | METRN | FCMR |
| C6orf62 | DNMT1 | DHX15 | STIM1 |
| USP9X | CHD3 | PFDN6 | PIGS |
| ACBD6 | WASHC3 | CYB5A | CYTH4 |
| DCAF15 | UTP18 | C1orf123 | IKBKE |
| ARHGAP17 | PIAS2 | CAPN10 | PQLC3 |
| ARHGEF1 | PLCB2 | TSC2 | MTSS1 |
| IRF2BP2 | ZNF224 | PCYT1A | ALDH9A1 |
| SENP6 | LRRC8A | MRPL44 | GPALPP1 |
| RBX1 | CHST11 | STN1 | TIGIT |
| BTG3 | ARMCX3 | ODC1 | ARL6IP1 |
| MRPS15 | TRPM7 | ZCCHC10 | PTGDR |
| DNAJC21 | PSENEN | PARP10 | RBM6 |
| PPA1 | RTN3 | PRKD3 | GID8 |
| RNF145 | PSTPIP1 | TCEAL3 | IRF2 |
| LSM12 | GLCCI1 | DSTYK | ZFAND2B |
| MSL3 | TWF1 | SLC52A2 | TRIM73 |
| CCDC57 | ATF7IP2 | MAN2A2 | KDM1A |
| MED17 | BTF3L4 | NOL4L | POM121 |
| TDG | ATXN2L | ADARB1 | POLR3K |
| UFL1 | SELL | FBXO6 | C7orf26 |
| CHCHD7 | AL118516.1 | SGK3 | AC074032.1 |
| NCBP2 | YIPF5 | SNX29 | CCDC32 |
| MAPRE2 | NUTM2A.AS1 | TUBGCP6 | PSMD10 |
| TAB2 | ARL5A | EMC10 | NCAPD3 |
| SLF1 | CD226 | LMO4 | ZSWIM8 |
| LDLRAP1 | TNPO1 | PDHB | OSBPL5 |
| HSPB11 | TESC | TPD52 | GON7 |
| GMCL1 | DCTN1 | ZBTB4 | SAFB |
| FLT3LG | FAM168B | MED29 | CMTM3 |
| CBL | HACD2 | GTF3C1 | HOTAIRM1 |
| RAMMET | IARS | RPL7L1 | DOK2 |
| AEBP2 | MRPS2 | RASGRP1 | PECAM1 |
| TNRC6C | THYN1 | SPG11 | KLRD1 |
| TXK | FAM53B | RCAN3 | LY9 |
| PMF1 | LTB4R | MAPRE1 | L3MBTL3 |
| PRKCQ | NIP7 | EPC2 | WDTC1 |
| MRNIP | CYB561 | RIPK1 | ZNF708 |
| MRPS11 | DAZAP1 | ST6GALNAC6 | ANAPC4 |
| OLA1 | MTMR6 | TSSC4 | SBNO1 |
| TSC22D2 | SAP30 | TRAF3 | ZNF91 |
| PRH1 | ZNF644 | SMARCD2 | CARHSP1 |
| DPP8 | SLC16A7 | SREBF2 | SEPT11 |
| ATF2 | ZFAND3 | EBAG9 | MIGA1 |
| SLC25A45 | SNRNP200 | CINP | R3HDM1 |
| KIAA0355 | PPP1R14B | NPEPL1 | COPB2 |
| DCK | BSG | RAD50 | RNF8 |
| TSPAN14 | AKR1A1 | SESN1 | TAB3 |
| ENTR1 | SLC38A2 | TSPAN31 | PDGFD |
| MIB1 | CD40LG | OAZ2 | SMAGP |
| CTSO | MAP2K6 | UBL7 | CLCN7 |
| RHOH | RGS16 | RARA | HCFC1R1 |
| NFKBIZ | TAF5L | RCC1 | SBK1 |
| NUCB2 | RNF20 | DPH5 | AMZ2 |
| GRK6 | UMPS | TET3 | HEATR1 |

*Continued on next page*

Supplementary Table 5 – *Continued from previous page*

| naive DN | CM DN | EM DN | EMRA DN |
|---|---|---|---|
| XPA | AC097376.2 | PCNX3 | CLASP2 |
| DGKA | KATNA1 | CIDEB | PLEKHA8 |
| PUF60 | SLC10A3 | NR1D1 | TRPT1 |
| SAMD1 | FUBP3 | AC009309.1 | AL365361.1 |
| TUBA1A | GFER | NCR3 | VIPR2 |
| BAZ1A | ASPSCR1 | ERP44 | METTL15 |
| G3BP2 | PEX13 | EMC4 | PPFIA1 |
| VHL | RRS1 | HEXIM1 | TSEN34 |
| VBP1 | DOPEY2 | JAML | CSK |
| SF3A3 | EAF2 | MRPL2 | CCL3L1 |
| ERO1B | PEX6 | SRPRB | CEMIP2 |
| CCDC137 | RHNO1 | DEGS1 | GPR174 |
| MRPS25 | TGIF2 | ZBTB16 | CYTOR |
| ARMCX6 | TRIM33 | LIN7C | ZC3H8 |
| TGIF1 | LINC00667 | PUS10 | RAP2A |
| ARRDC2 | FAM89B | KAT2B | SRPK2 |
| ITGA6 | RPA1 | SLC39A8 | PRKCB |
| NOL8 | PSMD1 | SLC12A7 | IKZF2 |
| BBIP1 | C12orf29 | DDX41 | ZFAND6 |
| DTX3L | PPP3CC | AGPAT4 | AC004687.1 |
| GCH1 | MGAT4B | TRIM24 | POLR2J3.1 |
| RFFL | CD5 | SORBS3 | UBL7.AS1 |
| RGS14 | KHSRP | ELP3 | CTPS1 |
| TXNDC15 | ALG2 | SLC25A22 | ELK4 |
| TMEM173 | TNFRSF14.AS1 | FOXJ2 | XCL2 |
| CHMP7 | DDRGK1 | GALK2 | PPTC7 |
| NSMCE2 | BRI3 | RTTN | SCRN2 |
| ZNF506 | CRTC2 | CARD8.AS1 | ADNP |
| PRKD2 | EFCAB2 | VRK3 | MEI1 |
| VPS13B | GALNT3 | GUCD1 | TTC13 |
| GORASP2 | COLQ | VAMP3 | AC098850.3 |
| EXOC3 | AC116366.3 | ITM2C | TMUB2 |
| FANCF | AUTS2 | RYBP | YLPM1 |
| LTBP3 | SPTSSA | GPR171 | CEP290 |
| CLTC | DNAJC17 | FIGNL1 | TARDBP |
| ZNF217 | KIAA0100 | VPS16 | ACTR3 |
| ZFAND1 | AARSD1 | SIK1 | STXBP3 |
| AC025164.1 | HOXB2 | CASP9 | LINC00944 |
| DICER1 | ANKRD40 | BBS7 | CAPN1 |
| AC087190.1 | SNAPC2 | ADAM19 | TERF1 |
| TRIM4 | GCDH | LRRC75A | NUMA1 |
| CDK5RAP2 | SASS6 | SUGP1 | ATP6AP1 |
| TFCP2 | RHOU | ACO2 | AP3D1 |
| CNOT10 | DHRS12 | SPTY2D1 | MUC20.OT1 |
| NUDT4 | GAS6.AS1 | CCDC91 | OXSR1 |
| PIK3C3 | EDEM2 | ASCC3 | MLLT10 |
| SCAF4 | AC013264.1 | TOB1 | SPPL2A |
| OTUD6B | CENPH | USP36 | MDM4 |
| THUMPD2 | AC132192.2 | GSR | MAPK1 |
| MPLKIP | BBS4 | SH3GLB1 | STT3B |
| MARS | SMAD7 | ILF2 | AP1S2 |
| CNOT11 | IRF9 | FUT11 | TMEM131 |
| IPO5 | NUP54 | PPIF | RYK |
| HDGFL3 | TMEM238 | GMPR2 | BPGM |
| AMMECR1 | RCN2 | SKIL | CS |
| CYB561A3 | CERK | TRIM69 | ZADH2 |
| TMEM218 | NOP10 | KIF5C | TOX |
| PI4K2A | YTHDF1 | TMEM63A | CENPK |
| ZNF580 | BCCIP | COG3 | NLRC3 |
| AC005261.1 | ZNF226 | IL18R1 | ZNF267 |
| MRFAP1L1 | KAT8 | NBEAL2 | CTDP1 |
| CIZ1 | RIC8A | NELFA | DGLUCY |
| GAN | AK2 | ARL4A | BBX |
| TTC7A | CUL3 | COG2 | SMAD2 |
| RAD17 | KLRC1 | THUMPD3 | PTK2B |
| SC5D | C16orf54 | ECPAS | ITGAL |
| PITPNA.AS1 | NINJ1 | JKAMP | PSMD11 |
| TCEA3 | SYTL2 | S100PBP | APOBEC3C |
| HELQ | RTKN2 | DCAF10 | CRTAM |
| PELP1 | SLC39A1 | RAE1 | RAB5A |
| MAGEF1 | ARL5B | PLXNA3 | OPTN |
| APBB1 | E2F3 | Z98885.3 | HNRNPH2 |

*Continued on next page*

Supplementary Table 5 – *Continued from previous page*

| naive DN | CM DN | EM DN | EMRA DN |
|---|---|---|---|
| ZMAT5 | MEGF9 | FUBP1 | ZFYVE16 |
| LMO7 | AQP3 | KMT5B | AP2A1 |
| METTL3 | SAE1 | CTSA | ZBTB38 |
| RRP1 | PPP1R12C | CAMK2D | TUT7 |
| COL18A1 | NKAPD1 | KIF20B | S1PR5 |
| RSRC1 | HPGD | EZH2 | DCP2 |
| ETNK1 | MRPL21 | ZNF791 | HLA.DRA |
| POLR2F | SIKE1 | DPP4 | PEF1 |
| PTCD3 | DCAF16 | VCAN | CLIP4 |
| LASP1 | OSBPL3 | POLR1E | NXPE3 |
| MORC3 | CD59 | ZNF746 | TRIM41 |
| KIN | C22orf46 | AEN | SMIM8 |
| PDE3B | ADAT1 | SNTA1 | CD244 |
| MAPKAPK5.AS1 | TTL | UBXN6 | VTA1 |
| MHENCR | HDAC8 | PRMT9 | PRSS23 |
| FAM208B | EPS8L2 | ZFYVE28 | VPS33A |
| SLC35A3 | MUTYH | GOLGA5 | CCL3 |
| NISCH | GCHFR | CEBPD | HAUS1 |
| ZNF439 | SCCPDH | LST1 | USP24 |
| NELL2 | MIR22HG | | DTHD1 |
| TRIM44 | AHCYL1 | | TCHP |
| AC008105.3 | GALM | | TSC22D1 |
| GRB2 | FBXO22 | | FCRL6 |
| ZNF131 | TRMT6 | | SP3 |
| PDE7A | | | ZMYND11 |
| TMEM131L | | | HLA.DQA2 |
| MNT | | | CLSTN1 |
| AC073332.1 | | | KDM5A |
| PGD | | | MRPS6 |
| MRTO4 | | | NCKAP1L |
| ZBTB40 | | | LINC01089 |
| GDAP2 | | | GABARAP |
| PDE4DIP | | | ZNF431 |
| FLAD1 | | | FCGR3A |
| NBAS | | | EP300 |
| CEP68 | | | ATG7 |
| TGFBRAP1 | | | FLNB |
| SLC11A1 | | | JAZF1 |
| CASP6 | | | DCUN1D5 |
| MFSD8 | | | FOPNL |
| INPP4B | | | ZNF236 |
| DIMT1 | | | LRRC47 |
| GNPDA1 | | | MTIF2 |
| TMEM70 | | | MASTL |
| DGAT1 | | | ORMDL2 |
| MTAP | | | FAM102B |
| SLC44A1 | | | TARS2 |
| ASB6 | | | CTBP2 |
| NUP160 | | | C1RL.AS1 |
| PRDX3 | | | ME2 |
| ARL1 | | | SENP5 |
| MLXIP | | | CUL4A |
| MRM3 | | | HSH2D |
| C18orf21 | | | RRAGC |
| TIMM21 | | | PAM |
| IFT52 | | | PCNX1 |
| BCL3 | | | NOM1 |
| NAB1 | | | ABHD2 |
| NEDD9 | | | ZEB2 |
| TBP | | | MOAP1 |
| ZSCAN18 | | | ORAOV1 |
| TMEM250 | | | SSFA2 |
| SUPT6H | | | MTURN |
| PINK1 | | | PMS1 |
| ASH1L.AS1 | | | C12orf43 |
| MTHFD2 | | | RIN3 |
| MFSD6 | | | DENND2D |
| SNX18 | | | PICALM |
| RXRB | | | ANKRD36C |
| SGK1 | | | LAIR2 |
| C1GALT1C1 | | | PRF1 |
| ZFP91 | | | GRK5 |

*Continued on next page*

Supplementary Table 5 – *Continued from previous page*

| naive DN | CM DN | EM DN | EMRA DN |
|---|---|---|---|
| CLPB | | | SERPINB6 |
| XRRA1 | | | KPNA5 |
| PARG | | | KLRF1 |
| WDR11 | | | AOAH |
| PCED1B | | | IL17RA |
| NUP58 | | | MRPS18C |
| PSMB5 | | | RNF219 |
| AL121603.2 | | | MED15 |
| SNN | | | BATF |
| LRRC37B | | | GNPAT |
| CNP | | | SNRNP35 |
| MBTD1 | | | TTC38 |
| SRP68 | | | DNM2 |
| INO80C | | | NOL9 |
| OSBPL2 | | | HDAC10 |
| MRI1 | | | CEP78 |
| ATP13A1 | | | NAPG |
| PLD3 | | | ZNF480 |
| ZNF329 | | | KIAA0586 |
| SPECC1L | | | FAM126B |
| RABL2B | | | CD160 |
| GTF3C2 | | | FABP5 |
| IMPACT | | | ADGRG1 |
| UPF1 | | | HIST1H2BN |
| DARS2 | | | CAMK2G |
| AC012360.3 | | | GZMH |
| CCDC58 | | | SACS |
| PRKCI | | | TRGV9 |
| MCCC1 | | | SESTD1 |
| SETD7 | | | ARID3B |
| AL121944.1 | | | |
| SKIV2L | | | |
| ORC3 | | | |
| CYP3A5 | | | |
| SMIM30 | | | |
| TIMM8A | | | |
| HMBS | | | |
| HSPA14 | | | |
| TIMM23 | | | |
| BICD1 | | | |
| NTHL1 | | | |
| CMTR2 | | | |
| LYRM9 | | | |
| COASY | | | |
| PSMG1 | | | |
| ID3 | | | |
| DCLRE1C | | | |
| MRPL58 | | | |
| MLLT11 | | | |
| YOD1 | | | |
| WDPCP | | | |
| RBM43 | | | |
| DIS3L2 | | | |
| RNF123 | | | |
| BRPF3 | | | |
| ZNF273 | | | |
| EPHX2 | | | |
| PVT1 | | | |
| SCAI | | | |
| EXOSC2 | | | |
| KCNC1 | | | |
| PLBD1 | | | |
| CRAMP1 | | | |
| RANBP10 | | | |
| HSBP1 | | | |
| ZNF606 | | | |
| ZNF208 | | | |
| PADI4 | | | |
| IL6R | | | |
| TTC24 | | | |
| MRPL53 | | | |
| USP40 | | | |

*Continued on next page*

Supplementary Table 5 – *Continued from previous page*

| naive DN | CM DN | EM DN | EMRA DN |
|---|---|---|---|
| SERPINI1 | | | |
| RHOBTB3 | | | |
| IRAK1BP1 | | | |
| CYSLTR1 | | | |
| FAM122C | | | |
| PKIA | | | |
| AC009812.1 | | | |
| SAMD12 | | | |
| MSRB2 | | | |
| AC004148.2 | | | |
| RPTOR | | | |
| DNAJB7 | | | |
| HIST2H2BE | | | |
| CR1 | | | |
| TCTE3 | | | |
| IFT74 | | | |
| VWA5A | | | |
| AC005842.1 | | | |
| TNFAIP2 | | | |
| SDK2 | | | |
| AC011445.2 | | | |
| ZNF134 | | | |
| SOCS3 | | | |
| THUMPD1 | | | |
| CCR7 | | | |
| DDX1 | | | |
| JAGN1 | | | |
| NPDC1 | | | |
| SERINC5 | | | |
| CREB1 | | | |
| SCML4 | | | |
| C15orf40 | | | |
| SF3B3 | | | |
| NSL1 | | | |
| DRAM2 | | | |
| PIGP | | | |
| USP11 | | | |
| TMEM179B | | | |
| KPNA3 | | | |
| ENOSF1 | | | |
| FAM76A | | | |
| TRAK2 | | | |
| FASTKD5 | | | |
| AHCTF1 | | | |
| GIPC1 | | | |
| BCL10 | | | |
| GPR155 | | | |
| NUPL2 | | | |
| MSH6 | | | |
| HPF1 | | | |
| FBXL22 | | | |
| RNF157 | | | |
| PASK | | | |
| PCNX4 | | | |
| TPP2 | | | |
| TESPA1 | | | |
| CSNK1E | | | |
| C1orf162 | | | |
| MYC | | | |
| EXOC6 | | | |
| ZNHIT3 | | | |
| SPRTN | | | |
| TCP11L2 | | | |
| ITGB3BP | | | |
| ABRAXAS1 | | | |
| FBXL3 | | | |
| RBM23 | | | |
| LEF1 | | | |
| APBA2 | | | |
| TRABD2A | | | |
| SETD3 | | | |
| CEP170 | | | |

*Continued on next page*

Supplementary Table 5 – *Continued from previous page*

| naive DN | CM DN | EM DN | EMRA DN |
|---|---|---|---|
| AP1B1 | | | |
| TMEM154 | | | |
| PNPLA6 | | | |
| NOL10 | | | |
| AC090948.1 | | | |
| GPR82 | | | |
| ZNF107 | | | |
| MAL | | | |
| EHBP1 | | | |
| SLC12A9 | | | |
| LHPP | | | |
| RNF43 | | | |
| AC062029.1 | | | |
| NME4 | | | |
| ZNF414 | | | |
| DERA | | | |
| ZNF486 | | | |
| EPHA1.AS1 | | | |
| BCL7A | | | |
| CACNA1I | | | |
| ROCK2 | | | |

**Supplementary Table 5.** Genes used in identification of DN step maturation

| naive DP | CM DP | EMRA DP |
|---|---|---|
| BOD1L1 | SMC5 | TAF15 |
| ELF2 | METTL26 | CLK1 |
| SELL | GLIPR1 | TRIP12 |
| PDE3B | SORL1 | ELP2 |
| GADD45B | SNRPF | M6PR |
| ZNF131 | CEBPZ | KIF21B |
| NAE1 | CFLAR | CTSD |
| GLS | KMT2A | ATP5MF |
| NUCB2 | EIF4G2 | BRD4 |
| THOC2 | IL2RG | BLOC1S1 |
| LDLRAP1 | METAP2 | MACF1 |
| ANKIB1 | CD247 | INPP5D |
| CALCOCO1 | NUDT22 | SOD2 |
| SRPRA | TFRC | TBCB |
| C15orf40 | IQGAP2 | MTPN |
| TRABD2A | IDH2 | KIF5B |
| NOSIP | PHAX | CCDC12 |
| HIST1H4C | ARPC4 | RNH1 |
| STIM2 | CA5B | TRIR |
| CLTB | TES | PDIA3 |
| CCDC91 | UBN1 | DNTTIP2 |
| GBP3 | TRMT112 | RNASET2 |
| ASB1 | SIAH2 | DUSP1 |
| AP3B1 | RAD23A | SAMHD1 |
| ZNF92 | NAP1L4 | TLE4 |
| AKAP17A | CAND1 | CDV3 |
| TRIM14 | DGKZ | SF3B5 |
| NMT2 | C12orf75 | HNRNPAB |
| TRIP11 | U2SURP | DDOST |
| MED15 | ORAI1 | CNP |
| LEF1 | SH3KBP1 | PAK1 |
| TAF1D | BCAP31 | CLCN7 |
| RALGAPA1 | RB1CC1 | GLUL |
| CCR7 | TAGAP | PRRC2A |
| PTCD3 | TGFB1 | PDZD8 |
| ZNF655 | SUPT6H | CDC42EP3 |
| CCT4 | MRPS35 | SETD2 |
| RABL6 | KLHDC3 | EIF2AK1 |
| GRPEL1 | RNF145 | CYC1 |
| FAM208A | GLOD4 | NCBP3 |
| RANBP9 | HDGF | FAM160B1 |
| GNAQ | POLR2F | HAUS3 |
| SURF4 | USP7 | GTF3C6 |
| ERC1 | LY9 | SCO1 |

*Continued on next page*

Supplementary Table 6 – *Continued from previous page*

| naive DP | CM DP | EMRA DP |
|---|---|---|
| CYB5B | BAZ1B | FAM210B |
| ZZEF1 | TOX4 | SELENOI |
| ZNF331 | MFNG | POLR3D |
| C8orf33 | HSPB11 | ABR |
| SCFD1 | ARHGEF6 | TUBA1A |
| RBMS1 | DEDD2 | SRGN |
| CCDC88B | TOMM22 | AP5Z1 |
| | WDFY1 | TXNRD1 |
| | BBIP1 | SMARCE1 |
| | DDX23 | GAB3 |
| | ABTB1 | NPRL3 |
| | DDRGK1 | RER1 |
| | VMAC | RILPL2 |
| | SYTL1 | NDUFB5 |
| | SSBP4 | SLC16A7 |
| | IL10RA | MED17 |
| | DHRS7 | NDUFA10 |
| | DOK2 | LYST |
| | ANXA1 | FAM126B |
| | ITM2A | TCERG1 |
| | CEBPZOS | RAB3GAP2 |
| | CERS2 | FGR |
| | MYDGF | SH3BP1 |
| | UPF3A | CD300A |
| | OXLD1 | MON1B |
| | PLEKHJ1 | CALM2 |
| | CTNNBL1 | IFT57 |
| | EXOG | FERMT3 |
| | UBE2Z | MAGOH |
| | ERN1 | NDUFC2 |
| | LATS1 | PSMD8 |
| | EMB | CTR9 |
| | NCBP2.AS2 | RIF1 |
| | SNX17 | MKNK2 |
| | COPB1 | REL |
| | SERGEF | ARF1 |
| | DEXI | NDUFB7 |
| | CRTC3 | PIP4K2A |
| | PIEZO1 | MRPS25 |
| | CBX1 | CWC27 |
| | UNC13D | CPNE1 |
| | RBMXL1 | THEMIS2 |
| | PYHIN1 | CX3CR1 |
| | SMIM15 | TMEM242 |
| | GTF2H5 | EEA1 |
| | SMS | DCAF4 |
| | HSD17B10 | SNX1 |
| | SMPD1 | FAM192A |
| | IMMP1L | POLDIP2 |
| | ARHGAP1 | PIGB |
| | ELMSAN1 | ZNRD1 |
| | C15orf61 | HEXA |
| | CCDC97 | PHC2 |
| | DERL1 | LIMS1 |
| | C5orf24 | ATG7 |
| | ITGB1 | BNIP1 |
| | PHYKPL | KDM7A |
| | GPATCH2L | CISD3 |
| | STAT4 | NR1H2 |
| | BCL2L11 | FBXO42 |
| | EPC2 | BTF3L4 |
| | MTRF1L | BCL10 |
| | NUPL2 | SCAMP3 |
| | SH2D1A | TMEM131 |
| | LLPH | CWC22 |
| | FAM32A | CYB561D2 |
| | ZNF224 | NEPRO |
| | AC073111.5 | LRPAP1 |
| | SMARCC2 | ARHGAP4 |
| | TMEM256 | TMEM109 |
| | MLEC | AP002387.2 |
| | LRRC41 | ATG16L2 |

*Continued on next page*

Supplementary Table 6 – *Continued from previous page*

| naive DP | CM DP | EMRA DP |
|---|---|---|
| | KYAT3 | STAM |
| | ACP6 | TSPAN31 |
| | SUSD4 | NHLRC3 |
| | TP53BP2 | HSBP1 |
| | TAF5L | CLTC |
| | SCCPDH | SYNGR2 |
| | MCFD2 | GIPC1 |
| | PRADC1 | EID2B |
| | DUSP28 | PHKG2 |
| | GLB1 | MOB4 |
| | UMPS | PTPN23 |
| | DNAJB11 | ARF4 |
| | NOA1 | REEP5 |
| | FBXO4 | FAM8A1 |
| | RNF14 | TENM1 |
| | HIST1H2BE | IDE |
| | SLC35B2 | PIK3AP1 |
| | PM20D2 | VCPKMT |
| | NME8 | SAMD4A |
| | TRGV2 | FAM219B |
| | GCC1 | PFDN4 |
| | TOX | CLASRP |
| | SGK3 | ARHGAP35 |
| | MFSD3 | KLF11 |
| | INVS | LINC00893 |
| | RDX | MRPL13 |
| | PTS | GTF3C5 |
| | REXO2 | CORO1C |
| | ROBO3 | NAXD |
| | CREM | DNAJA3 |
| | ZNF33B | AC005921.2 |
| | PYROXD2 | REXO1 |
| | PHC1 | KLF12 |
| | PLEKHA5 | RHOC |
| | ZNF641 | WDR45B |
| | RXYLT1 | EPS15 |
| | PITPNM2 | ADGRE5 |
| | NCOR2 | ARIH2 |
| | LINC02361 | PYCARD |
| | FBXO34 | MIS18BP1 |
| | FAM96A | TMBIM4 |
| | TMEM8A | MOB1A |
| | KCTD13 | IER5 |
| | ELP5 | PLEK |
| | SHMT1 | VASP |
| | GHDC | STMP1 |
| | TSEN54 | MRPS16 |
| | RAB40B | PDP1 |
| | AP000845.1 | MRM3 |
| | TP53INP2 | C1orf123 |
| | TPD52L2 | ABT1 |
| | ADAMTS10 | AC114760.2 |
| | MRI1 | CNOT8 |
| | GEMIN7 | METTL2B |
| | CRKL | NDUFA9 |
| | IGLC2 | CCNK |
| | AL031846.2 | PLCB2 |
| | CRELD2 | YIPF2 |
| | SBF1 | PANK2 |
| | CFAP298 | ZNF217 |
| | DUSP2 | TIA1 |
| | APEX1 | MARCH8 |
| | INO80B | STAT5A |
| | MGAT5 | ELOF1 |
| | DCUN1D1 | ZKSCAN1 |
| | TRIM23 | MAPK6 |
| | DHX16 | ZBTB4 |
| | AKIRIN2 | ZSWIM7 |
| | TMEM168 | SARS |
| | GSPT2 | LIAS |
| | KIF22 | CCDC112 |
| | CBX4 | TUBGCP5 |

*Continued on next page*

Supplementary Table 6 – *Continued from previous page*

| naive DP | CM DP | EMRA DP |
|---|---|---|
| | RINL | CD79B |
| | DNAJC3 | FGFBP2 |
| | ZNF836 | LYZ |
| | | ZEB2 |
| | | PUSL1 |
| | | THAP3 |
| | | FBXO6 |
| | | PLOD1 |
| | | MFN2 |
| | | KDM1A |
| | | INPP5B |
| | | SF3A3 |
| | | SLC16A1 |
| | | RFX5 |
| | | ZBTB41 |
| | | AIDA |
| | | ZNF678 |
| | | RDH14 |
| | | MIR4435.2HG |
| | | POLR2D |
| | | PIKFYVE |
| | | LANCL1 |
| | | CNOT9 |
| | | CRELD1 |
| | | CTNNB1 |
| | | ZNF445 |
| | | UQCRC1 |
| | | KBTBD8 |
| | | MRPS22 |
| | | CCDC50 |
| | | GAK |
| | | NELFA |
| | | RCHY1 |
| | | ARFIP1 |
| | | MARCH1 |
| | | TMEM267 |
| | | PPWD1 |
| | | MAST4 |
| | | DMXL1 |
| | | LINC01184 |
| | | ISOC1 |
| | | MED7 |
| | | MXD3 |
| | | NUP153 |
| | | E2F3 |
| | | SLC39A7 |
| | | AL451165.2 |
| | | PPP2R5D |
| | | TMEM14A |
| | | AGPAT4 |
| | | YAE1D1 |
| | | TAF6 |
| | | UBN2 |
| | | CHPF2 |
| | | INSIG1 |
| | | PDK3 |
| | | CASK |
| | | UTP14A |
| | | ELF4 |
| | | RBMX2 |
| | | SLC9A6 |
| | | TNKS |
| | | AC108863.1 |
| | | TRMT10B |
| | | SPTLC1 |
| | | HABP4 |
| | | COQ4 |
| | | MRPS2 |
| | | RNF141 |
| | | CAT |
| | | TRAF6 |
| | | TIMM10 |

*Continued on next page*

Supplementary Table 6 – *Continued from previous page*

| naive DP | CM DP | EMRA DP |
|---|---|---|
| | | SYVN1 |
| | | FUT4 |
| | | ENDOD1 |
| | | SIDT2 |
| | | OTUD1 |
| | | ARHGAP21 |
| | | CSTF2T |
| | | EIF4EBP2 |
| | | P4HA1 |
| | | RRP12 |
| | | CHUK |
| | | STN1 |
| | | COPS7A |
| | | CMAS |
| | | DCTN2 |
| | | ATP2B1.AS1 |
| | | UHRF1BP1L |
| | | FBXW8 |
| | | DDX51 |
| | | PGAM5 |
| | | NUP58 |
| | | TEP1 |
| | | AL136295.5 |
| | | PPP2R3C |
| | | TXNDC16 |
| | | SYNE3 |
| | | TRMT61A |
| | | PACS2 |
| | | NIPA2 |
| | | GCHFR |
| | | GALK2 |
| | | ZNF280D |
| | | ANKDD1A |
| | | NPTN |
| | | HMG20A |
| | | C16orf91 |
| | | HCFC1R1 |
| | | GLYR1 |
| | | GDE1 |
| | | EEF2K |
| | | SETD1A |
| | | ITGAM |
| | | LYRM9 |
| | | WIPF2 |
| | | NARF |
| | | NPC1 |
| | | PLCB1 |
| | | SLC9A8 |
| | | CAPS |
| | | PNPLA6 |
| | | PRAM1 |
| | | ATG4D |
| | | DNASE2 |
| | | LPAR2 |
| | | TIMM50 |
| | | PPP5C |
| | | ALDH16A1 |
| | | PTOV1 |
| | | JOSD2 |
| | | ZNF880 |
| | | SLC25A1 |
| | | APOL2 |
| | | RANGAP1 |
| | | TRPM2 |
| | | AL031280.1 |
| | | EYA3 |
| | | MYCBP |
| | | EVI5 |
| | | LRIG2 |
| | | NBPF14 |
| | | SRGAP2 |
| | | BOLA3.AS1 |

*Continued on next page*

Supplementary Table 6 – *Continued from previous page*

| naive DP | CM DP | EMRA DP |
|---|---|---|
| | | LINC01943 |
| | | GPBAR1 |
| | | ARL8B |
| | | RPUSD3 |
| | | ZNF619 |
| | | TCAIM |
| | | KLHL18 |
| | | RFT1 |
| | | AC093010.2 |
| | | POGLUT1 |
| | | HES1 |
| | | NRROS |
| | | CTBP1 |
| | | AC147067.1 |
| | | KIAA0232 |
| | | CENPU |
| | | F2R |
| | | FAM172A |
| | | AC116366.1 |
| | | HSPA4 |
| | | TGFBI |
| | | PPP2R2B |
| | | CSF1R |
| | | EXOC2 |
| | | PAK1IP1 |
| | | GFOD1 |
| | | KIF13A |
| | | SOX4 |
| | | DAXX |
| | | FGD2 |
| | | PEX6 |
| | | NDUFAF4 |
| | | DCBLD1 |
| | | SMPDL3A |
| | | FUCA2 |
| | | FIGNL1 |
| | | SLC12A9 |
| | | PIK3CG |
| | | TMEM140 |
| | | ACOT9 |
| | | CXorf38 |
| | | CDK16 |
| | | GNL3L |
| | | ARHGEF9 |
| | | UPRT |
| | | AIFM1 |
| | | DNASE1L1 |
| | | AGPAT5 |
| | | AF131216.1 |
| | | OTUD6B |
| | | AC103706.1 |
| | | CNTLN |
| | | ZBTB5 |
| | | ABCA1 |
| | | ELP1 |
| | | PTRH1 |
| | | FUBP3 |
| | | SLC2A6 |
| | | CHID1 |
| | | AC132192.2 |
| | | ADM |
| | | KAT5 |
| | | EIF1AD |
| | | RIN1 |
| | | DCUN1D5 |
| | | PDGFD |
| | | SDHD |
| | | ADO |
| | | IFIT2 |
| | | MMS19 |
| | | FAM45A |
| | | STK32C |

*Continued on next page*

Supplementary Table 6 – *Continued from previous page*

| naive DP | CM DP | EMRA DP |
|---|---|---|
| | | NINJ2 |
| | | DCP1B |
| | | LPCAT3 |
| | | CLEC12A |
| | | DDX47 |
| | | EPS8 |
| | | AC092747.4 |
| | | IPO8 |
| | | OAS3 |
| | | CAMKK2 |
| | | SLC7A7 |
| | | ESR2 |
| | | TMEM63C |
| | | TRAF3 |
| | | CKB |
| | | PARP6 |
| | | FURIN |
| | | CLEC16A |
| | | CETP |
| | | CIAPIN1 |
| | | COQ9 |
| | | CLEC18A |
| | | PDPR |
| | | CMTR2 |
| | | ZCCHC14 |
| | | TAX1BP3 |
| | | ATAD5 |
| | | ADAP2 |
| | | MYO19 |
| | | AC060780.1 |
| | | G6PC3 |
| | | AC005332.7 |
| | | CD300C |
| | | CD300E |
| | | SLC26A11 |
| | | SLC16A3 |
| | | EPB41L3 |
| | | SLC39A6 |
| | | SLC23A2 |
| | | ABHD12 |
| | | NECAB3 |
| | | EDEM2 |
| | | MAFB |
| | | GNA15 |
| | | ICAM4 |
| | | PDE4A |
| | | MAP1S |
| | | PROSER3 |
| | | ZNF574 |
| | | CD33 |
| | | SIGLEC10 |
| | | YPEL1 |
| | | CSF2RB |
| | | CD63 |
| | | MYO1G |
| | | IQSEC1 |
| | | TENT5A |
| | | SEC16A |
| | | CCL4L2 |
| | | ECPAS |
| | | PER1 |
| | | LILRB1 |
| | | TKT |
| | | BID |
| | | FKBP1A |
| | | LINC01934 |
| | | CTSZ |
| | | TCF7L2 |
| | | RRAGC |
| | | SLC50A1 |
| | | MEF2D |
| | | SMC6 |

*Continued on next page*

Supplementary Table 6 – *Continued from previous page*

| naive DP | CM DP | EMRA DP |
|---|---|---|
| | | BABAM2 |
| | | PNPT1 |
| | | MRPS9 |
| | | TMEM87B |
| | | DZIP3 |
| | | WDR55 |
| | | HLA.DQA2 |
| | | MEA1 |
| | | RAB32 |
| | | CYBB |
| | | PCSK1N |
| | | CHM |
| | | THAP1 |
| | | CHCHD7 |
| | | SLC15A3 |
| | | SDHAF2 |
| | | OS9 |
| | | APPL2 |
| | | SIPA1L1 |
| | | PSEN1 |
| | | ZNF410 |
| | | TMED8 |
| | | SPATA33 |
| | | MIR22HG |
| | | CRLF3 |
| | | RAB31 |
| | | PET117 |
| | | AC245060.5 |
| | | SRGAP2B |
| | | G0S2 |
| | | AC108463.3 |
| | | CAMK1 |
| | | EAF2 |
| | | MEF2C |
| | | HLA.DQA1 |
| | | HLA.DQB1 |
| | | LFNG |
| | | EIF4EBP1 |
| | | ALDH3B1 |
| | | SLC39A9 |
| | | ITGAX |
| | | COG1 |
| | | CD300LF |
| | | CEBPA |
| | | PLXNB2 |
| | | RAB37 |
| | | CEBPB |
| | | ARID3A |
| | | IER5L |
| | | CFD |
| | | FCGR3A |
| | | ZDBF2 |
| | | POLD3 |
| | | PILRA |
| | | CFP |
| | | AP2A1 |
| | | HMOX1 |
| | | HES4 |
| | | NCF2 |
| | | LMO2 |
| | | SMIM25 |

**Supplementary Table 6.** Genes used in identification of DP step maturation
